## Supplementary figures and images for "Human dynein-dynactin is a fast processive motor in living cells"

### Figure 1 - figure supplement 1

#### Figure 1 - figure supplement 1

A

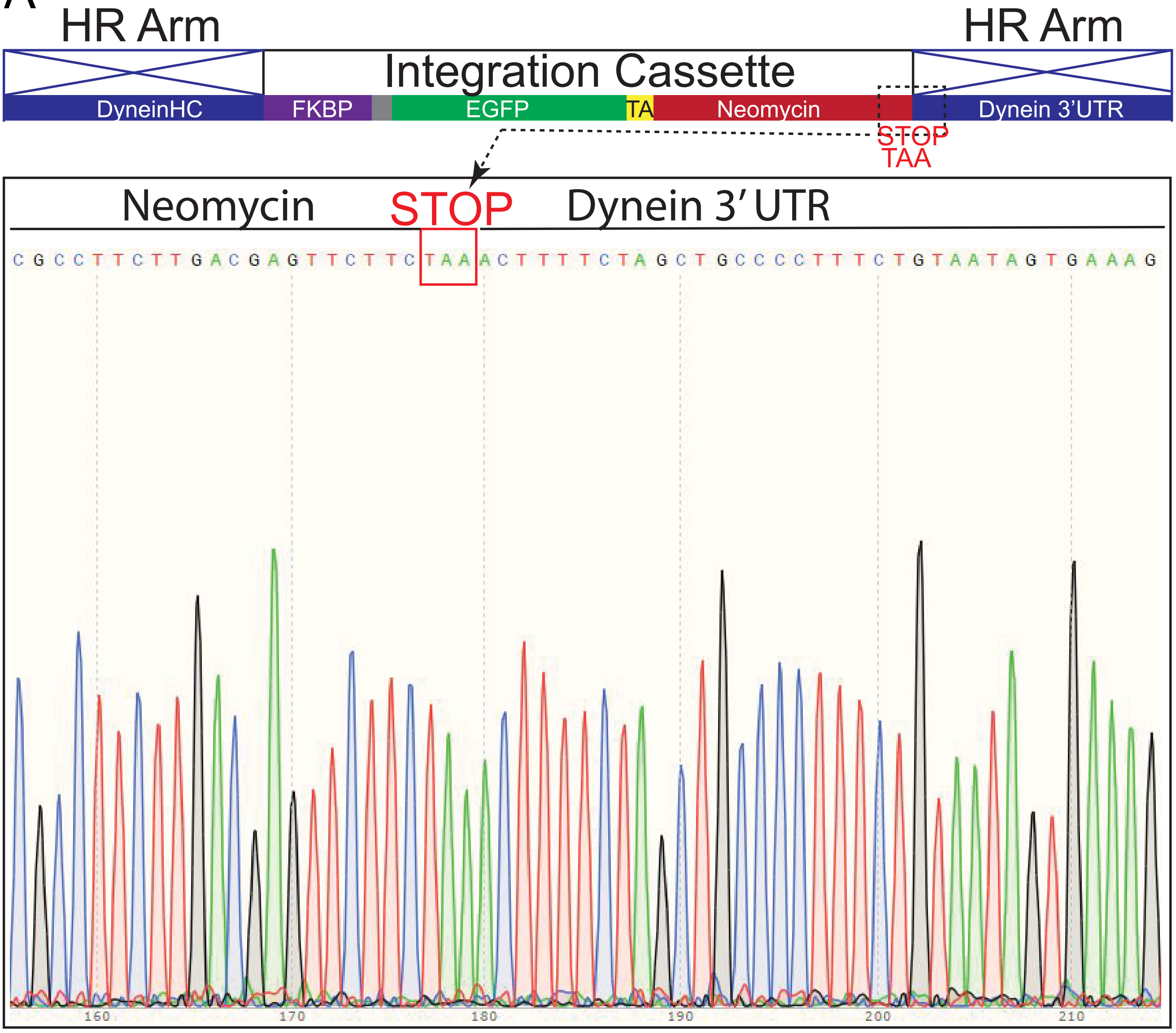


B

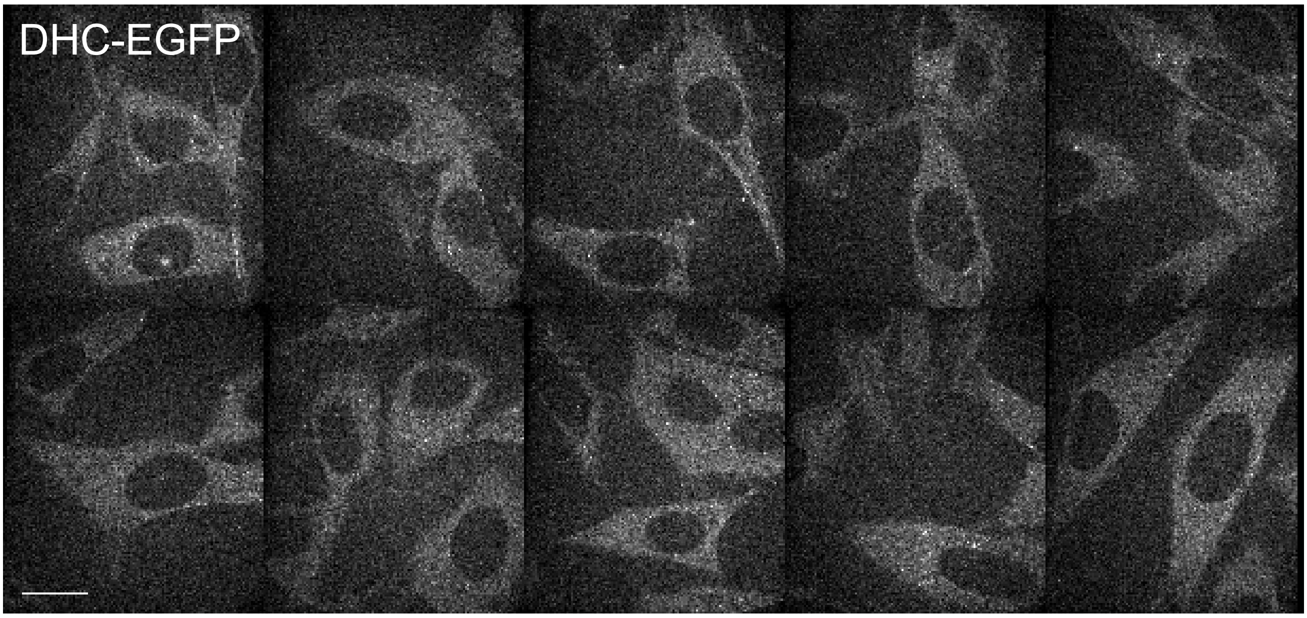

### Figure 2 - figure supplement 1

#### Figure 2 - figure supplement 1

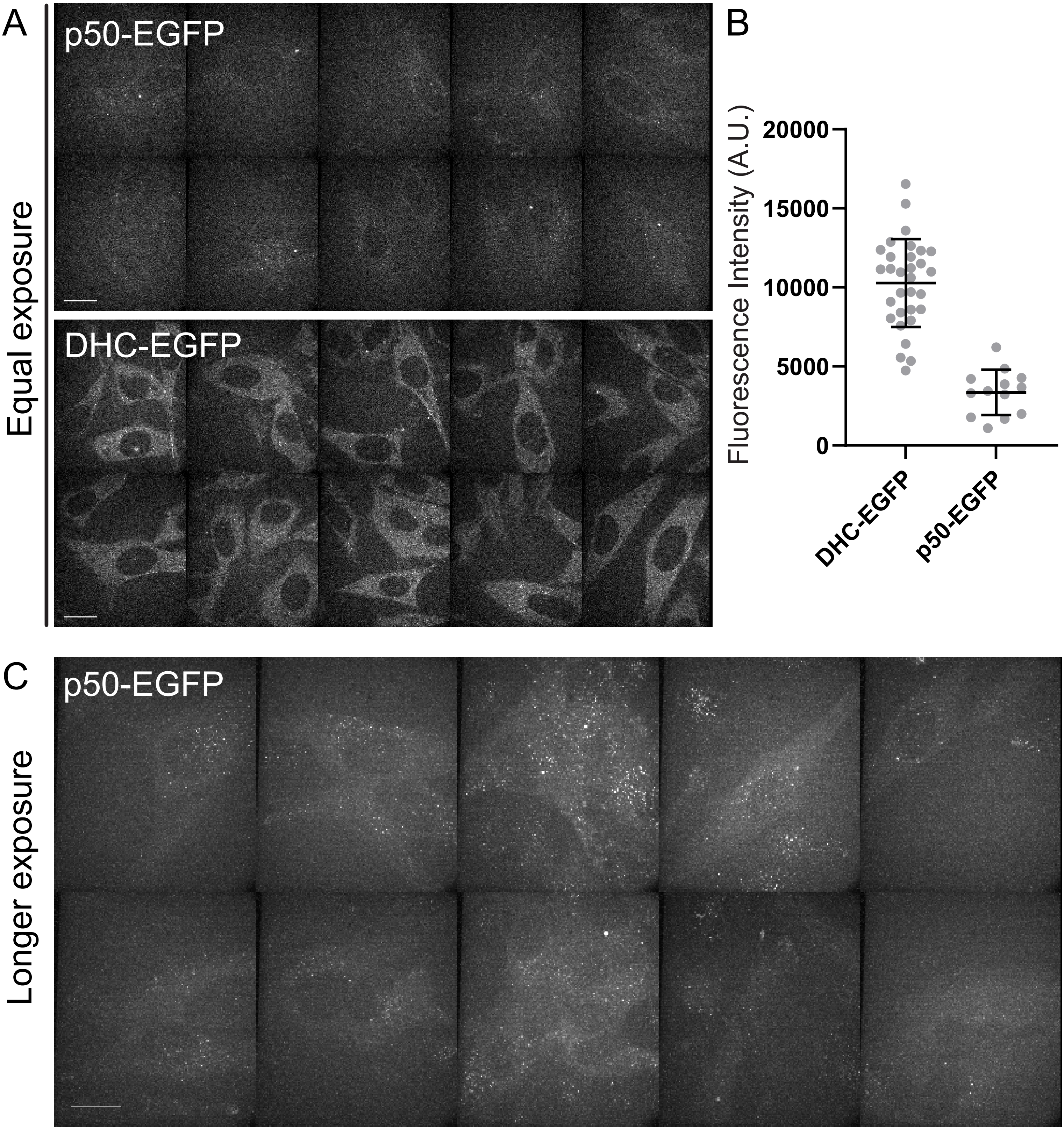

### Figure 3 - figure supplement 1

Figure 3 - figure supplement 1

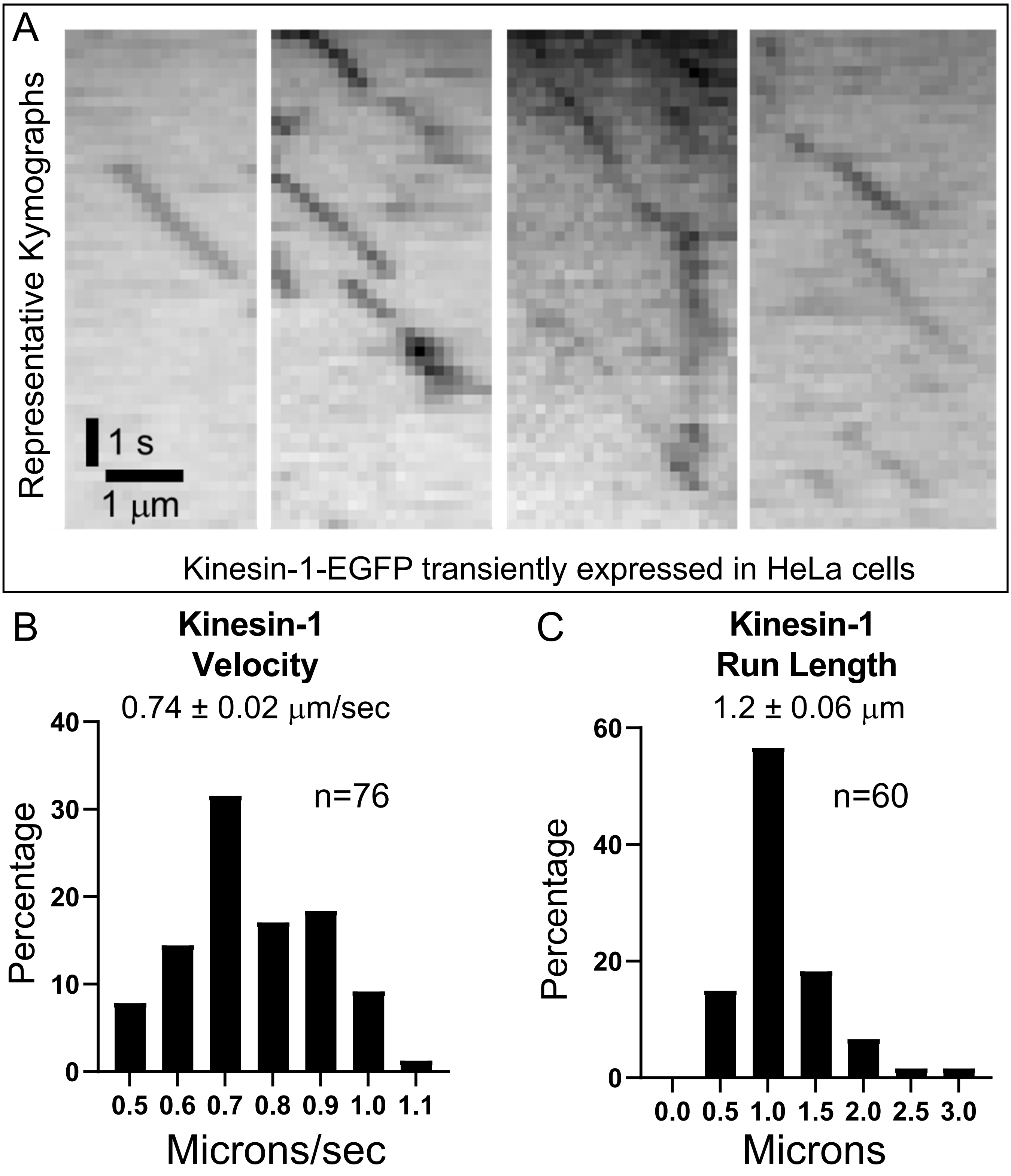

### Figure 3 - figure supplement 2

#### Figure 3 - figure supplement 2

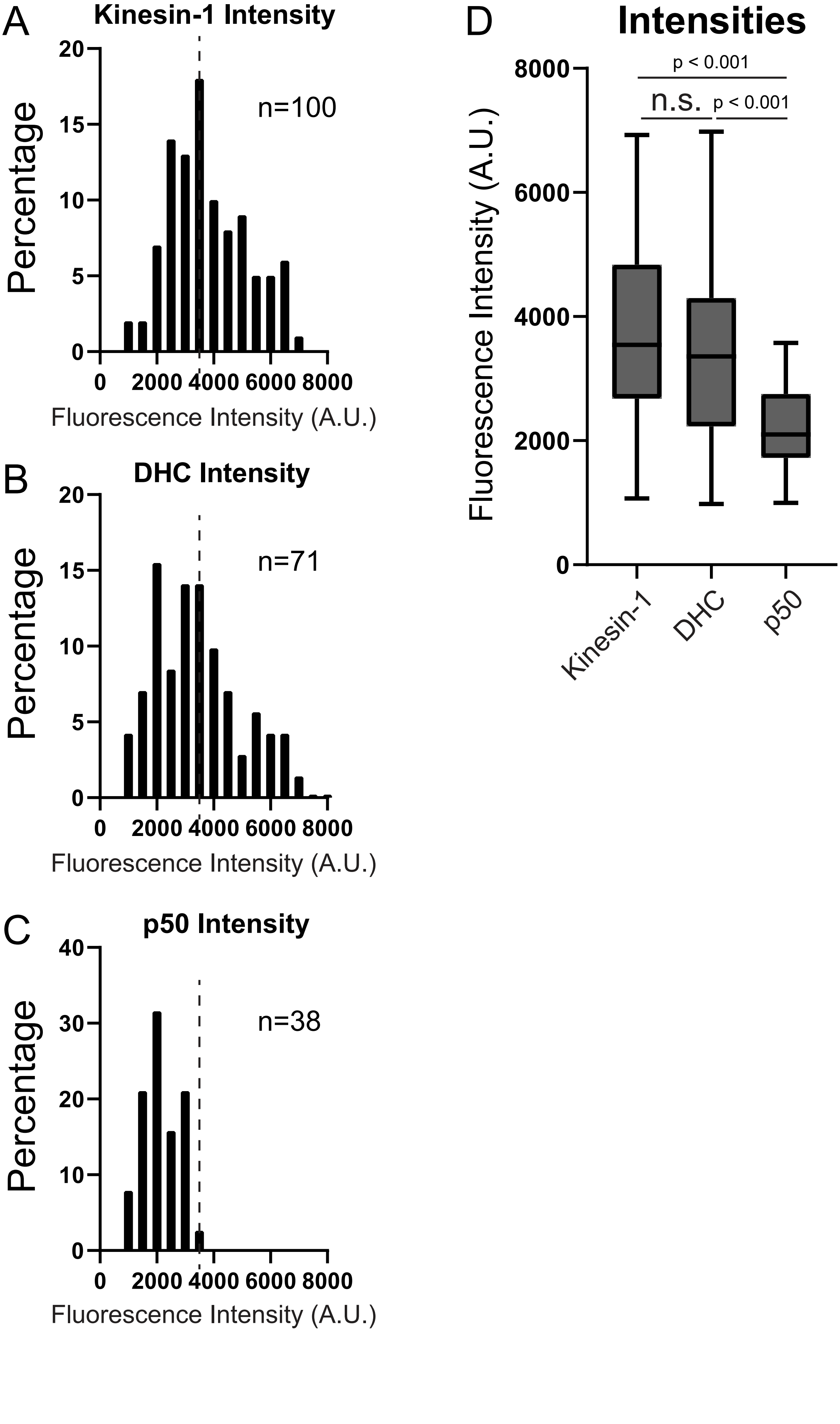
